# Global lysine acetylation in *Escherichia coli* results from growth conditions that favor acetate fermentation

**DOI:** 10.1101/457929

**Authors:** Birgit Schilling, Nathan Basisty, David G. Christensen, Dylan Sorensen, James S. Orr, Alan J. Wolfe, Christopher V. Rao

**Author notes:** To whom correspondence should be addressed: Christopher V. Rao, Department of Chemical and Biomolecular Engineering, University of Illinois at Urbana-Champaign, Urbana, Illinois, 61801, USA. Alan J. Wolfe, Department of Microbiology and Immunology, Stritch School of Medicine, Health Sciences Division, Loyola University Chicago, CTRE 224, 2160 South First Avenue, Maywood, Illinois, 60153, USA.

## Abstract

Lysine acetylation is thought to provide a mechanism for regulating metabolism in diverse bacteria. Indeed, many studies have shown that the majority of enzymes involved in central metabolism are acetylated and that acetylation can alter enzyme activity. However, the details regarding this regulatory mechanism are still unclear, specifically with regards to the signals that induce lysine acetylation. To better understand this global regulatory mechanism, we profiled changes in lysine acetylation during growth of *Escherichia coli* on the hexose glucose or the pentose xylose at both high and low sugar concentrations using label-free mass spectrometry. The goal was to see whether lysine acetylation differed during growth on these two different sugars. No significant differences, however, were observed. Rather, the initial sugar concentration was the principal factor governing changes in lysine acetylation, with higher sugar concentrations causing more acetylation. These results suggest that acetylation does not target specific metabolic pathways but rather simply targets accessible lysines, which may or may not alter enzyme activity. They further suggest that lysine acetylation principally results from conditions that favor accumulation of acetyl phosphate, the principal acetate donor in *E. coli*.

**IMPORTANCE:** Bacteria alter their metabolism in response to nutrient availability, growth conditions, and environmental stresses. This process is best understood at the level of transcriptional regulation, where many metabolic genes are conditionally expressed in response to diverse cues. However, additional modes of regulations are known to exist. One is lysine acetylation, a post-translational modification known to target many metabolic enzymes. However, unlike transcriptional regulation, little is known about this regulatory mode. We investigated the factors inducing changes in lysine acetylation by comparing growth on glucose and xylose. We found that the specific sugar used for growth did not alter the pattern of acetylation; rather, the principal factor was the amount of sugar, with more sugar yielding more acetylation. These results imply lysine acetylation is a global regulatory mechanism that is not responsive to the specific carbon source per se but rather the accumulation of downstream metabolites.

## OBSERVATION

N^ε^-lysine acetylation is an abundant post-translational modification in many bacteria (1, 2). Multiple studies have shown that lysine acetylation predominantly targets the enzymes involved in central metabolism (2–5). Because these lysines are often catalytically active, their acetylation may regulate metabolism in bacteria. This hypothesis is supported by a handful of *in vitro* studies showing that lysine acetylation indeed alters the activity of some enzymes involved in central metabolism (4, 6–8). However, it is still not clear what role lysine acetylation plays in regulating metabolism. One hypothesis is that lysine acetylation provides a global mechanism by which cells regulate metabolism in response to their energy status. The response to the energy status occurs through the availability of the acetyl-group donors: acetyl-CoA and acetyl phosphate, two metabolic intermediates that are related through a single reaction. According to this model (2), lysine acetylation reduces carbon flux through central metabolism when this flux exceeds the capacity of the tricarboxylic acid (TCA) cycle, for example by acetate overflow metabolism (9).

We sought to explore this hypothesis by profiling changes in lysine acetylation during growth of *Escherichia coli* on the hexose D-glucose or the pentose D-xylose. Our goal was to see whether lysine acetylation differed during growth on these two different sugars. We chose to compare the effects of these two sugars because *(i) E. coli* grows faster on glucose than on xylose and thus may have different lysine acetylation patterns (10); *(ii)* their metabolism involves a number of different enzymes, which may also be differentially acetylated; and *(iii)* these are the two most abundant sugars in plant biomass (11). Due to the relevance for biomass production, *E. coli* has been genetically engineered to produce a wide variety of valuable chemicals and fuels from these sugars, with the goal of replacing petroleum-based feedstocks with renewable, plant-based ones (12, 13). These designs require that carbon flux is redirected towards producing these compounds. Thus, any knowledge regarding the regulation of carbon flux during growth on these two sugars will aid in these engineering efforts.

To measure changes in lysine acetylation, cells from three biological replicates were harvested after 12 hours of growth in M9 minimal medium (**Figure S1A and S2** and then subjected to label-free mass spectrometry (data-independent acquisition, DIA), as described previously (14, 15), for details also see **Text S1**. Protein expression was similar under the four different growth conditions, with only 61 proteins identified exhibiting significant changes in expression (**Figure S3**). Many differentially expressed proteins were less abundant during growth at high sugar concentrations. Reduced expression likely results from catabolite repression due to high sugar concentrations (**Table S1**), as many of these proteins are expressed from CRP-dependent promoters (16). Fewer proteins exhibited increased expression during growth at high sugar concentrations. Four (AmtB, RutA, GlnK and Nac) are members of the NtrC (NRI) regulon, which is induced when the carbon-to-nitrogen ratio is high (17). Two (ProV and ProX) are components of the glycine betaine/proline ABC transporter, which is induced in response to osmotic stress (18). These results are consistent with high sugar concentrations inducing osmotic stress and being sensed by the cell as nitrogen limitation.

Label-free mass spectrometry identified 3840 unique acetyllysine sites on 978 proteins (**Tables S2-3**). Of these, 278 lysines on 157 unique proteins exhibited significant changes in acetylation (Figure 1A; **Table S4**). The relative degree of acetylation increased at least two-fold for 260 sites on 149 proteins during growth on 4% versus 0.4% glucose. No sites were found to exhibit a two-fold decrease in the relative degree of acetylation at the higher sugar concentration. Similarly, the relative abundance of acetylation increased at least two-fold for 256 sites on 147 proteins during growth on 4% versus 0.4% xylose. Once again, no sites exhibited a two-fold decrease in the relative abundance of acetylation at the higher sugar concentration.

**Figure 1.**
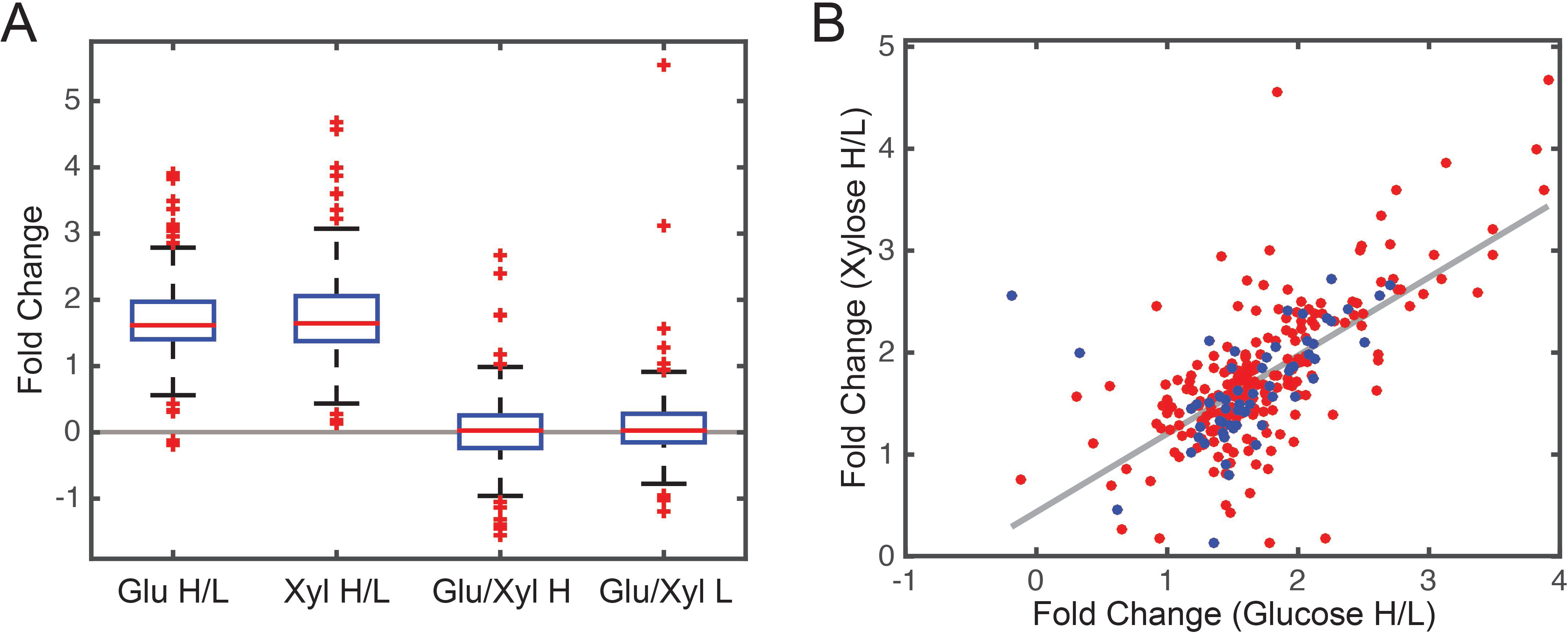
Relative changes in lysine acetylation under the four growth conditions. A. Box plot showing relative change in acetylation for the four different growth conditions. B. Comparison of differentially acetylated lysines during growth on xylose versus glucose. The blue dots denote lysines on the metabolic enzymes depicted in Figure 2. Abbreviations: Glu H/L (4% glucose versus 0.4% glucose); Xyl H/L (4% xylose versus 0.4% xylose); Glu/Xyl H (4% glucose versus 4% xylose); and Glu/Xyl L (0.4% glucose versus 0.4% xylose).

We next explored whether observed increases in the relative abundance of acetylation at higher sugar concentrations were correlated during growth on glucose versus xylose. As shown in Figure 1B, changes in acetylation are moderately correlated during growth on the two sugars (R^2^=0.45). In other words, many of those lysines where acetylation increased during growth on high glucose concentrations often increased during growth on high xylose concentrations. We separately confirmed these results using anti-acetyllysine western blots, which again showed that global acetylation is correlated with the concentration but not the identity of the carbon sources (**Figure S4**). These results suggest that many differentially acetylated lysines are not sensitive to the specific growth sugar but rather the amount of sugar available. They also support the hypothesis that most acetylation is a result of overflow metabolism and is independent of the specific route for catabolism. Indeed, cells grown on higher concentrations of sugar produced significantly more acetate (**Figure S1**).

We further examined those acetylated proteins exhibiting significant changes in their relative acetylation abundance, focusing on those involved in central metabolism (Figure 2). Only two of these enzymes exhibited significant (2-fold) changes amongst the four growth conditions: xylose isomerase (XylA) and phosphoenolpyruvate carboxylase (Ppc). XylA catalyzes the first step in xylose metabolism and is differentially acetylated at two sites. Interestingly, the relative abundance of acetylation for one lysine (XylA^K17^) increased, while the other (XylA^K381^) decreased during growth on 4% glucose versus 4% xylose. In fact, it is the only lysine on a metabolic enzyme to exhibit a decrease in the relative abundance of acetylation. The acetylation of lysine 17 also increased during growth on 4% versus 0.4% xylose, suggesting that it is sensitive to the energy state of the cells. These results imply that acetylation of these lysines may alter enzymatic activity. However, when we replaced these lysines with glutamines or arginines, which mimic acetylated lysine or unacetylated lysine, respectively (19–21), we observed no effect during growth on xylose (**Figure S5**). While these experiments may not accurately model lysine acetylation, they nonetheless suggest that acetylation is not regulating XylA activity. Little is known about the lysines for Ppc.

**Figure 2.**
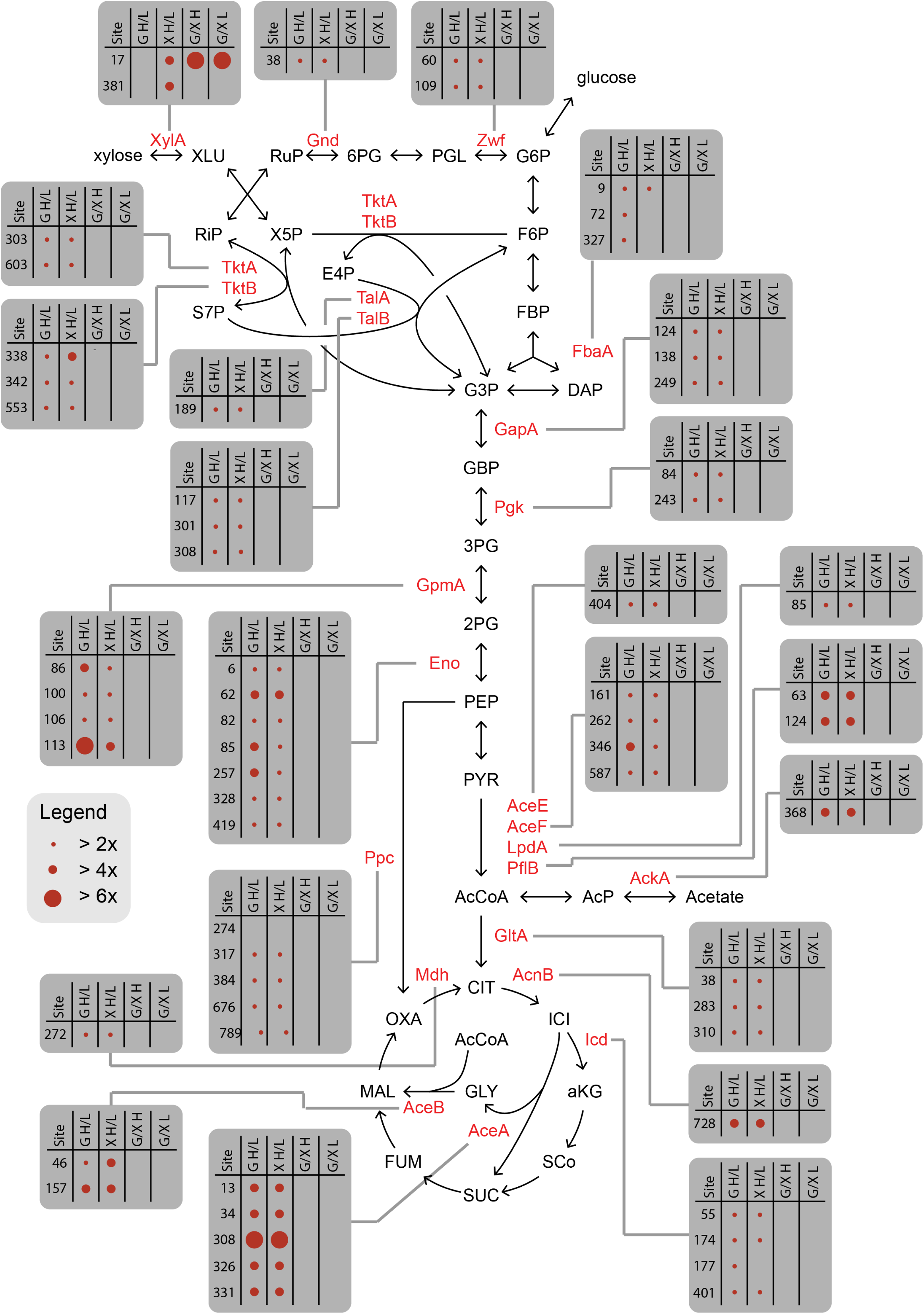
Enzymes in central metabolism exhibiting changes in lysine acetylation under the four growth conditions. Specific lysines are shown in gray boxes. Data is also available in **Table S3**. Abbreviations: XLU: xyluose; RuP: ribulose 5-phosphate; 6PG: gluconate 6-phosphate; PGL: phosphogluconolactone; G6P: glucose 6-phopshate; RiP: ribulose 5-phosphate; X5P: xylulose 5-phosphate; F6P: fructose 6-phosphate; S7P: sedoheptulose 7-phosphate; E4P: erythrose 5-phosphate; FBP: fructose 1,6-bisphosphate; G3P: glyceraldehyde 3-phosphate; DHAP: dihydroxyacetone phosphate; 3PG; GBP: 1,3-bisphosphoglycerate; 3PG: 3-phosphoglycerate; 2PG: 2-phosphoglycerate; PEP: phosphoenolpyruvate; PYR: pyruvate; AcCoA: acetyl-CoA: AcP: acetyl-phosphate; CIT: citrate; ICI: isocitrate; aKG: α-ketoglutarate; SCo: succinyl-CoA; SUC: succinate; FUM: fumarate; MAL: malate; OXA: oxaloacetate; Glu H/L (4% glucose versus 0.4% glucose); Xyl H/L (4% xylose versus 0.4% xylose); Glu/Xyl H (4% glucose versus 4% xylose); and Glu/Xyl L (0.4% glucose versus 0.4% xylose).

## Conclusion

We profiled protein acetylation in *E. coli* during growth on glucose and xylose at both high and low sugar concentrations. We did not observe major differences amongst the lysines acetylated during respective growth on these two sugars. Rather, the observed changes in lysine acetylation were principally correlated with the initial sugar concentration, with higher sugar concentrations causing more acetylation. These results indicate that acetylation is agnostic to the metabolic route and simply targets accessible lysines, which may or may not alter enzyme activity. They further support the hypothesis that lysine acetylation results from the buildup of metabolic intermediates, principally acetyl phosphate, under conditions that favor acetate production (**Figure S1**).

## ACKNOWLEDGMENT

This work was supported by an NCRR shared instrumentation grant for the TripleTOF 6600 (1S10 OD016281, Buck Institute) and by the U. S. Department of Energy, Office of Science, Office of Biological and Environmental Research under award number DE-SC0012443.

## SUPPLEMENTAL MATERIALS

**Text S1.** Supplemental materials and methods.

**Table S1.** Protein exhibiting significant changes in relative abundance.

**Table S2.** Mass spectrometric analysis of acetyl-enriched peptide fractions. Overall, a total of 3,840 unique acetylation sites and a total of 978 acetylated proteins were identified by tandem mass spectrometry (MS/MS).

**Table S3.** Identified acetylation sites during growth of *E. coli* on M9 minimal medium containing 0.4% glucose, 4% glucose, 0.4% xylose, or 4% xylose as the sole carbon source.

**Table S4.** Sites exhibiting significance changes in acetylation during growth of *E. coli* on M9 minimal medium containing 0.4% glucose, 4% glucose, 0.4% xylose, or 4% xylose as the sole carbon source.

**Figure S1.** Growth of *E. coli* on M9 minimal medium containing 0.4% glucose, 4% glucose, 0.4% xylose, or 4% xylose as the sole carbon source. Error bars denote the standard deviation for three biological replicates. A. Cell growth as determined by optical absorbance; B. sugar consumption during growth on 0.4% glucose or xylose; C. acetate production during growth on 0.4% glucose or xylose; D. sugar consumption during growth on 4% glucose or xylose; and E. acetate production during growth on 4% glucose or xylose. During growth on higher concentrations of sugar, consumption is incomplete due to acidification of the growth medium.

**Figure S2.** Mass spectrometric workflow to assess relative changes in acetylation site levels. Protein lysates are proteolytically digested and acetylated peptides are enriched by anti-acetyl affinity enrichment. Acetylated peptides are identified and quantified using modern proteomics technology (specifically data-independent acquisition, also referred to as SWATH).

**Figure S3.** Differential protein expression under the four growth conditions. Cells were harvested after 12 hours of growth. For mass spectrometric analysis, *E. coli* cells were grown as described above in M9 minimal media supplemented with i) 0.4% glucose, ii) 4% glucose, iii) 0.4% xylose, or iv) 4% xylose. Isolated frozen bacterial pellets from each of the 4 growth conditions (3 biological replicates each) were lysed, followed by tryptic digestion of the protein lysate and mass spectrometric acquisition of all samples by data independent acquisition. Protein expression values are provided in **Table S1**.

**Figure S4.** Relative changes in lysine acetylation under the four growth conditions as determined using anti-acetyllysine western blot.

**Figure S5.** Comparison of growth for different XylA mutants during growth in (A) 0.4% xylose or (B) 4% xylose.

